# Dashing 2: genomic sketching with multiplicities and locality-sensitive hashing

**DOI:** 10.1101/2022.10.16.512384

**Authors:** Daniel N. Baker, Ben Langmead

**Affiliations:** Department of Computer Science, Johns Hopkins University

## Abstract

A genomic sketch is a small, probabilistic representation of the set of k-mers in a sequencing dataset. Sketches are building blocks for large-scale analyses that consider similarities between many pairs of sequences or sequence collections. While existing tools can easily compare 10,000s of genomes, relevant datasets can reach millions of sequences and beyond. Popular tools also fail to consider *k*-mer multiplicities, making them less applicable in quantitative settings. We describe a method called Dashing 2 that builds on the SetSketch data structure. SetSketch is related to HyperLogLog, but discards use of leading zero count in favor of a truncated logarithm of adjustable base. Unlike HLL, SetSketch can perform multiplicity-aware sketching when combined with the ProbMinHash method. Dashing 2 integrates locality-sensitive hashing to scale all-pairs comparisons to millions of sequences. Dashing 2 is free, open source software available at https://github.com/dnbaker/dashing2

## 1 Introduction

Sketching, e.g. based on MinHash or HyperLogLog, is a key building block for scaling sequence comparison. Sketches built over all the k-mers in a sequence have been applied in clustering [1], phylogenetic inference [2], strain-level profiling [3, 4], species delineation [5] and summarization of genomic collections [6, 7]. While existing tools like Mash [1] and Dashing [7] can easily cluster 10,000s of genomes, many relevant biological datasets are much larger, reaching millions of sequences and beyond. Further, these tools fail to consider multiplicities of the k-mers, limiting their applicability in settings where quantities matter, e.g. when analyzing collections of sequence reads, or summaries from quantitative sequencing assays.

Dashing 2 builds on the recent SetSketch structure [8]. SetSketch is related to HyperLogLog (HLL), but replaces the HLL’s leading zero count (LZC) operation with a truncated logarithm of adjustable base. This addresses a major disadvantage of the HLL as implemented in Dashing, since the LZC wastes about 2 bits of space out of every 8-bit estimator (“register”) stored. SetSketch also has similarities to multiplicity-aware approaches like BagMinHash [9] and ProbMinHash [10]. All three approaches (SetSketch, BagMinHash and ProbMinHash) make decisions about whether and how to update registers by performing a random draw from a distribution, where the draw is seeded by a hash value derived from the input item. This allows SetSketch to perform multiplicity-aware sketching in the same way as the other sketches. SetSketch also admits a simple, accurate algorithm for computing similarity between sketches in a joint fashion.

Dashing 2 uses locality-sensitive hashing (LSH) to scale all-pairs comparisons to very large inputs. It finds near neighbors by grouping samples with equal register groupings. This makes Dashing 2 particularly effective for all-pairs comparisons over large sequence collections.

Dashing 2’s implementation of the SetSketch structure is efficient and versatile. Like the original Dashing software, Dashing 2 can be run in a mode that sketches a sequencing dataset and saves the result to a file. This is activated by the `dashing2 sketch` command. Also like Dashing, Dashing 2 can compare sequences or sketches in an all-pairs fashion (`dashing2 cmp` or, equivalently, `dashing2 dist`). When combined with the `--cache` option, Dashing 2 loads pre-existing sketches from disk, making the command much faster. When the input to these commands consists of many sketches or datasets, Dashing 2 performs all-pairs comparisons and outputs tabular results. Dashing 2’s new LSH-assisted all-pairs comparison mode can be activated via the `--similarity-threshold` *x* option, where e.g. *x* = 0.8 instructs Dashing 2 to use an LSH approach to consider only pairs whose similarity is likely to be 80% or higher.

Dashing 2 supports a range of sequence alphabets, including the 2-bit DNA alphabet and a standard 20-letter amino acid protein alphabet (using `--protein` option) and compressed amino acid alphabets of size 14, 8, and 6 (`--protein14`, `--protein8`, and `--protein6`) as described by [11]. Compressed protein alphabets are more appropriate when sequence identity is low.

Dashing 2’s new multiplicity-aware sketching mode can be enabled for sequencing data inputs via the `--prob` option. Dashing 2 can sketch BigWig inputs [12] encoding numerical coverage vectors using the `--bigwig` option. The more generic `--wsketch` mode can sketch inputs consisting of keys and weights.

Besides the modes discussed above, Dashing 2 has modes for computing containment coefficients, symmetric containment coefficient, and intersection size. Further, Dashing 2 has modes for computing Jaccard coefficients in an exact manner, without sketching or estimation; this is useful for evaluation but comes at the expense of longer running time and larger memory footprint compared to sketching-based approaches.

## 2 Methods

A sketching method distills a large dataset into a smaller collection of representative items. The MinHash method, for example, distills a dataset consisting of many items into a smaller set of just the *k* items that are *minimal*, i.e. having minimal hash values as computed by a hash function.

Since genomic data usually takes the form of a sequence collection, we must first convert such a collection to a mathematical set. This is typically accomplished by transforming the sequences into the set of their constituent length-*k* substrings, i.e. their *k*-mers. Since sequences that are reverse complements of each other should be considered identical, *k*-mers are usually canonicalized before being added to the set. That is, if a given *k*-mer is greater than its reverse complement, it is replaced with its reverse complement.

Though sketches are usually much smaller than the original dataset, they can still be used to estimate various relevant quantities, such as the cardinality of a dataset, i.e. how many distinct items/*k*-mers are present. Further, sketches for two different datasets can be used to estimate the similarity between the datasets. This can be done in far less memory or time compared to that required to compare the original, unsketched datasets.

### 2.1 SetSketch

Dashing [7] used the HyperLogLog data structure and its “leading zero count” (LZC) strategy to update register values. Specifically: an incoming item is hashed and its bitwise representation is partitioned into a prefix *p* and a suffix *q*. The prefix *p* determines which register the item maps to, while the suffix *q* determines how the register should be updated. Specifically, the algorithm finds the number of consecutive unset bits in the most significant digits of *q*, i.e. its LZC. If the LZC is greater than the register’s current value, the value is set to the LZC. The LZC serves as a kind of order-of-magnitude estimator; many such estimators can be averaged to accurately estimate cardinalities and similarities.

Whereas Dashing’s HLL registers were each 8 bits wide and able to hold a value in the range 0–255, LZCs could range only from 0 to 64. In fact, LZCs could would usually span a smaller range than this, since bits used for the prefix *p* are not considered. An LZC would therefore fail to use at least 2 bits of an 8-bit register, leaving the structure 25% empty. Though registers could be shrunk to 6 bits, this would conflict with Dashing’s use of SIMD instructions with 8-bit operands.

Ertl’s SetSketch [8] addresses this issue by replacing the LZC with a logarithm of configurable base *b*. This comes with a drawback: the addition of a single item to the SetSketch potentially updates the values of *all* registers, rather than just one. Given a data item *d*, the update rule for each register *K*_*i*_, is:

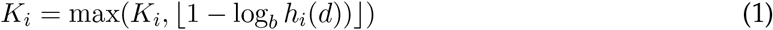

where *h*_*i*_ is an independent hash function specific to register *i* distributed exponentially; i.e. *h*_*i*_(*d*) ∼ Exp(*a*).

For reasons explained below, it is useful to factor this update rule into two phases, with delaying the logarithms to the second phase. We use *K*_*i*_ to denote the register’s value at update time and 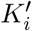 to denote its value after the final truncation.

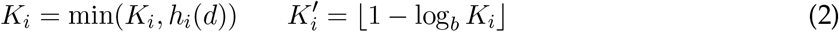

Where again *h*_*i*_(*d*) ∼ Exp(*a*). The subtraction in the 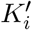 formula inverts the notion of “extremeness” from a minimum to a maximum, hence the use of min in Equation 2 versus the use of max in Equation 1.

The strategy of letting each register’s value be a function of all the input items has advantages. First: since the final value is a function of all items, rather than a register-specific subset of them, the final values are statistically independent. Second: the exponential rate *a* and logarithm base *b* are parameters of the sketch. We can set *b* in a way that spreads the *K*_*i*_ values over a range of our choosing. If registers are 8 bits wide, we can choose *a* and *b* so that ⌊1 − log_*b*_ *h*_*i*_(*d*))⌋ ranges from 0 to 255, landing outside the range only with low (and controllable) probability. If registers are another size (e.g. Dashing 2 supports 4, 8, 16, 32 and 64-bit registers), *a* and *b* can be adjusted. Methods for setting *a* and *b* are described in section 2.2 of [8].

This comes with a potential disadvantage. Since each register is a function of every input item, each addition may require *O*(*m*) work where *m* is the number of registers. Ertl [8] proposes optimizations that ensure that the work per update quickly becomes *O*(1) in the typical case where the input is much larger than *m*. This is accomplished by (a) maintaining a value *K*_max_ equal to the maximum among all the registers’ current values, and (b) reordering the inner loop so that iterations occur in increasing order by value of the *h*_*i*_(*d*) ∼ Exp(*a*) draw. Note that the maximum is maintained over the original, untruncated exponential draws in the *K*_*i*_ variables, not the truncated version eventually stored in the 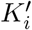 variables. Once we reach an iteration where the draw has value *h*_*i*_(*d*) *> K*_max_, neither the current nor a subsequent iteration can possibly change the value of a register, and the inner loop can break.

Another disadvantage of the SetSketch is the computational cost of the exponential draws and, related to that, the computational cost of logarithms. While the strategy just described reduces the number of exponential draws, later sections describe how we reduce the cost of logarithms.

### 2.2 Similarity comparison

Dashing 1’s default algorithm for estimating the Jaccard coefficient between HLLs *A* and *B, J* (*A, B*), used the Maximum Likelihood Estimation (MLE) method of [13]. This models register values as Poisson random variables and uses an iterative root-finding to estimate the Poisson parameter from the histogram of register values in the union (*A* ∪ *B*) HLL. This was less accurate but substantially faster than the related Joint MLE (JMLE) method [13], which required histogramming of joint register values.

Dashing 2 uses a simpler joint estimator named 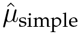 [8] ^1^. It has a closed-form solution and does not require an iterative root-finding procedure. Being a “joint” estimator, 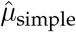 also does not require a union sketch; it is a function only of the input sketches *A* and *B*. Finally, 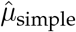 does not require a histogram of register *values*. Instead, it requires two counts: *D*_+_ and *D*_−_. *D*_+_ is the number of registers in *A* that are greater than their counterparts in *B* and *D*_−_ is the number of registers in *A* that are less than their counterparts. Unlike MLE and JMLE histograms, *D*_+_ and *D*_−_ can be computed using only Single-Instruction Multiple-Data (SIMD) instructions. In particular, a combination of SIMD greater-than/less-than and population-count instructions enable rapid tallying of *D*_+_ and *D*_−_ with respect to chunks of registers at a time. As described in [8], a mathematical complication arises when the sets are mostly disjoint. We fall back on Ertl’s alternative formulae *α*_disj_ and *β*_disj_ from [8] in such cases.

### 2.3 Full Dashing 2 sketch update

Algorithm 1 gives the update algorithm for the full version of the Dashing 2 SetSketch. This is the default for weighted sketching, and can be enabled for non-weighted sketching using the option `--full-setsketch`. Without this option, the one-permutation strategy described in the next subsection is used instead. Inputs consist of the register array *K* comprising the sketch, a MaxTree array *T* maintaining maxima over power-of-two-sized stretches of registers, the item *X* to be added, and the item’s weight *W*. When used for unweighted sketching, *W* equals 1. Registers in *K* are initialized to the maximum possible value.

A single update could require modifying any number of registers, from 0 to *m*. To avoid *O*(*m*) work on average, the algorithm follows the strategy of Ertl [8], examining registers in order according to the probability it will be updated, i.e. according to the extremeness that register’s exponential random draw. The algorithm uses a pseudo-random number generator RNG, seeded with *X* for deterministic updates. RNG.nextExponentialSpacing(*i, m, W*) returns the value that must be added to obtain the next draw in increasing order, according to the recurrence 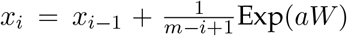. The recurrence follows from the memoryless property of the exponential distribution [10]. RNG.nextFisherYates() samples a new register randomly without replacement with Fisher-Yates shuffling, as in Algorithm 6 of [10]. By matching the increasing series of exponential draws with a random sequence of register choices, we visit registers in the desired order of most to least likely to be updated. Kahan summation [14, 15] is used to reduce numerical errors when summing exponential spacings.

Conditional statements on lines 4 and 12 of Algorithm 1 abort upon reaching a draw that is less extreme than the least extreme that could affect a register value (*K*_max_). Accordingly, the *K*_max_ ← T.update(*K*[*i*], *i*) statements on lines 9 and 17 update a structure maintaining the current “least extreme draw” in the *K*_max_ variable per Algorithm 4 of [10]. Updates to the *K*_max_ structure take *O*(log(*m*)) time, and are required only when a register is modified. The number of loop iterations is inversely related to the number of items added, approaching 0 as the number far exceeds *m*.

#### Algorithm 1

Update Full Dashing 2 SetSketch

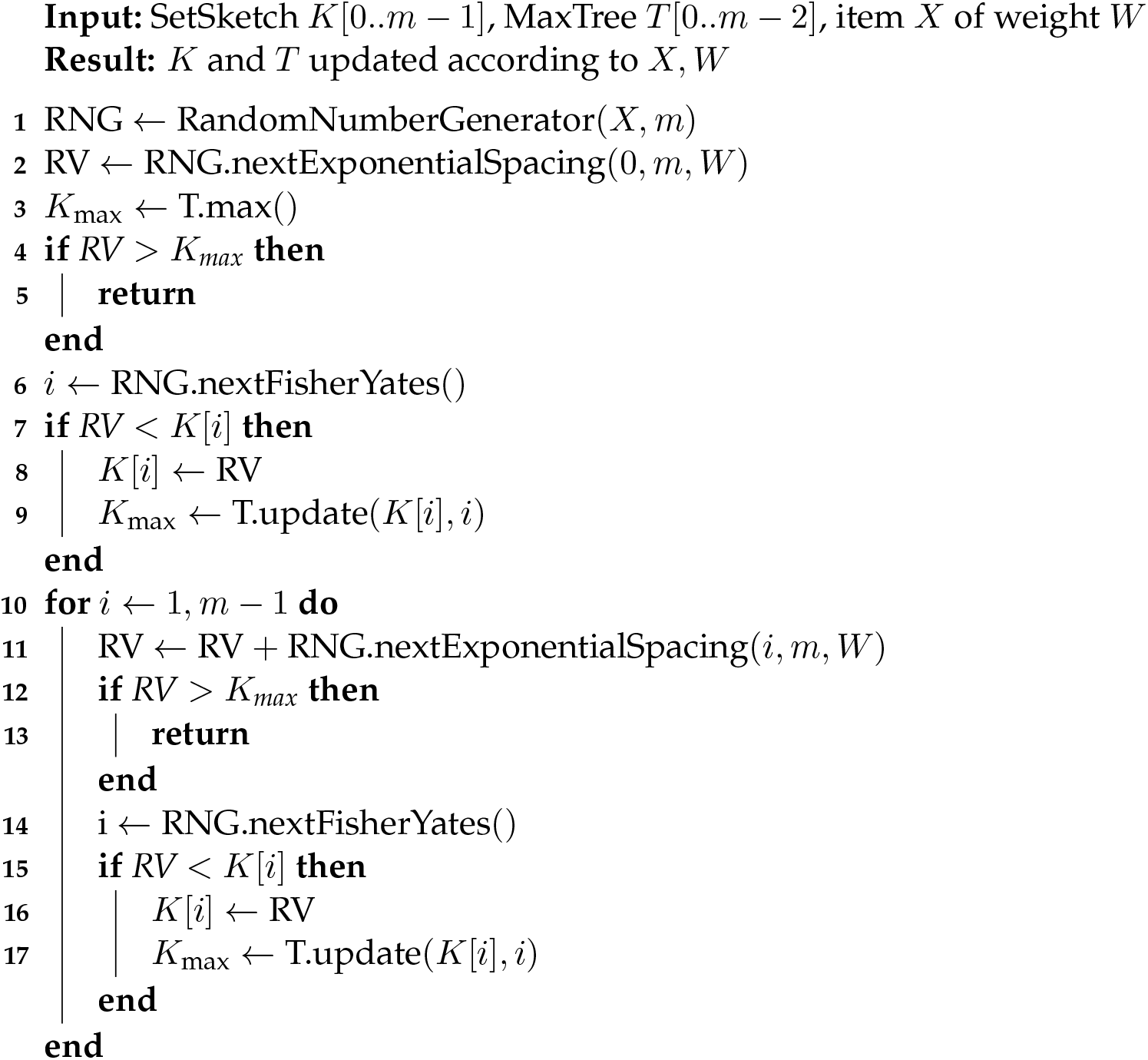

### 2.4 One-permutation SetSketch

By default, Dashing 2 computes unweighted sketches and uses an economical “one-permutation” update method [16], which modifies at most one register per update. This method uses some bits of the random draw to choose which register to update (Algorithm S2, Supplementary Material), similarly to how the HLL update rule in Dashing uses the hash prefix *p*.

While the one-permutation approach is efficient compared to a full update, accuracy suffers when many registers are empty, i.e. when the input has few items relative to *m*. To maintain accuracy, we implement the densification approach of [17], applied after finalization. This strategy has similar accuracy compared to the full SetSketch, but is more efficient in practice. Because of this, we made this one-permutation mode the default for unweighted sketching. The full update

#### Algorithm 2

Find candidates for *k* nearest neighbors for dataset (*K, id*) using LSH index. Result is a list of at most 3·*k* candidates that is later refined and ordered using Jaccard similarity. rule is used instead when the user enables weighted sketching, or when the user enables it with the `--full-setsketch` option.

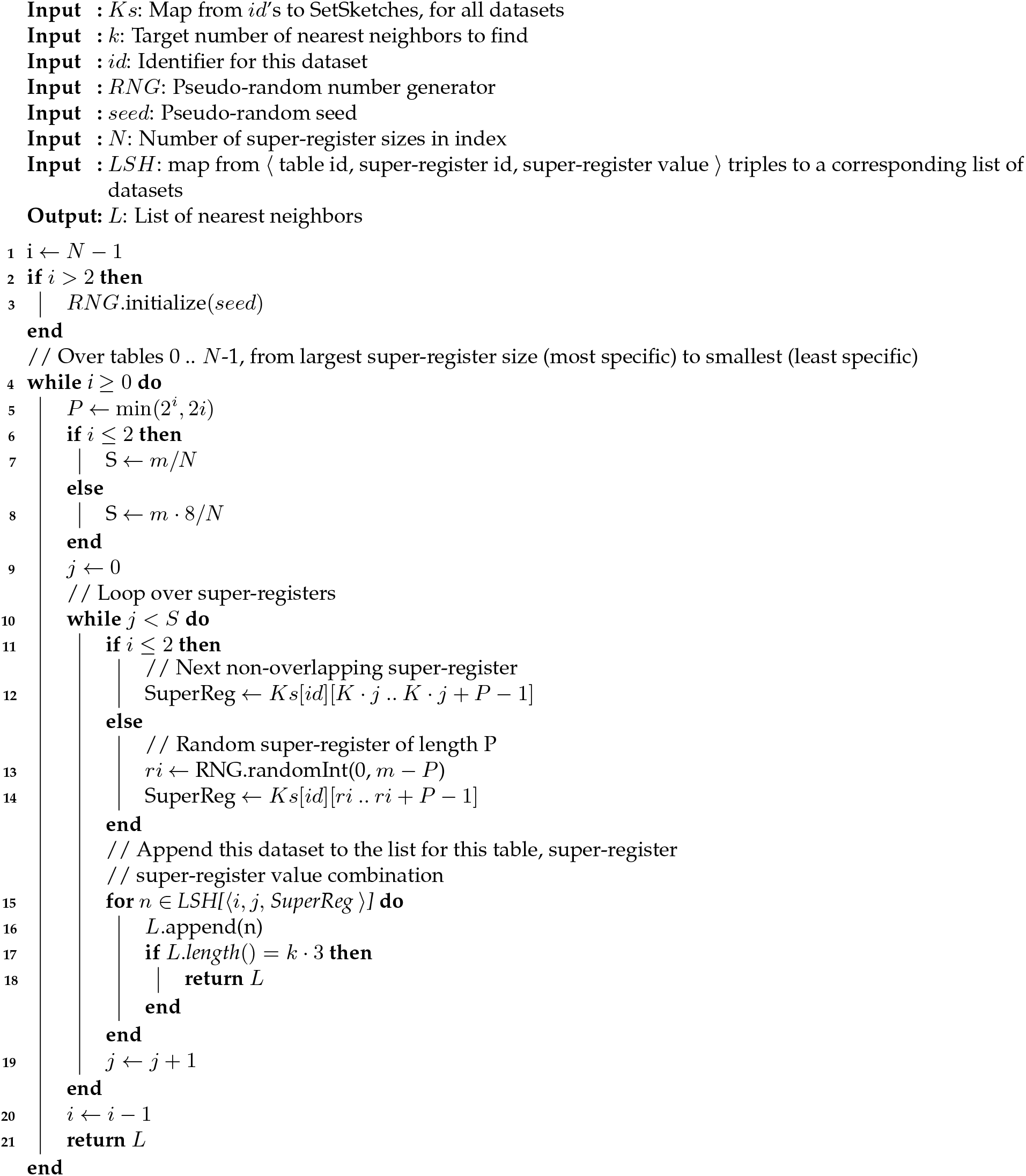

### 2.5 SetSketch parameters

Like the HyperLogLog, the number of registers *m* is a parameter of the SetSketch. Unlike Hyper-LogLog, SetSketch has a related parameter, the register *width* in bits. Other key parameters are the rate for the exponential draws (*a*), and the log base used for truncation needed to fit draws into registers (*b*). As in [8], we use *q* to denote the value that is 1 less than the maximum register value. E.g. for 8-bit registers, *q* = 2^8^ − 1 − 1 = 254.

While [8] gives theoretical guidelines for choosing *a, b* and other parameters, these assume foreknowledge of input cardinalities. On the other hand, multiple SetSketches are comparable only if they were built using identical parameters. This creates a tension between wishing to choose the parameters sooner, in order to make compact sketches, versus later, to delay truncation until we can ensure all relevant sketches are constructed and truncated with identical *a, b* and *q*.

Dashing 2’s default strategy is to set *a* and *b* according to the overall set of input datasets, and to shape and truncate the sketches according to user-configurable choices for *m* and *q*. Dashing 2 also allows the user to delay the choices for *a, b* and *q*, so that larger, untruncated sketches can be stored temporarily in preparation for future truncation and comparison with other sketches.

To select *a* and *b* according to the data, Dashing 2 first forms untruncated sketches, then computes *b* and *a* according to the expressions:

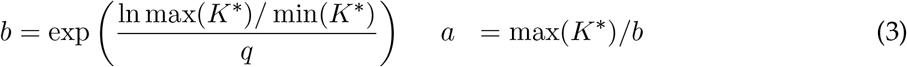

Where *K*^*^ denotes a concatenations of all untruncated register values from all inputs. For experiments in this study, we invoked Dashing 2 with all input datasets at once, ensuring *a* and *b* are set identically for all.

### 2.6 Delayed logarithms

Potentially expensive logarithm calculations are used in two tasks: (a) truncation of register values, and (b) to perform the exponential draws. Dashing 2 avoids these costs in two ways. First, it uses the two-step strategy of Equation 2 to delay truncation until a finalization step, which runs only after all items are added (Algorithm S3, Supplementary Material). Before finalization, intermediate register values are stored as un-truncated 64-bit floating-point numbers. As a result, the update rule uses only a minimum, rather than both a logarithm and a maximum. The total number of logarithmic truncations performed at most *m* regardless of the number of items added to the sketch.

This comes at the cost of requiring additional space at sketching time. For the entire algorithm up to finalization, we must store a 64-bit value for each register even if finalization will later reduce that to, e.g. 8 bits. Since practical sketches require only thousands of registers, this is not onerous in practice.

### 2.7 Approximate logarithms

The second use of logarithms is in the exponential Exp(*a*) random draw, which is accomplished by computing − ln(Unif())*/a* where Unif() is a uniform random draw between 0 and 1. We observed that, once the sketch becomes quite full, many exponential draws are well above the *K*_max_ ceiling, aborting the inner loop. While an inaccurate logarithm might cause us to miscompute whether a draw is under the *K*_max_ ceiling, this arises only for draws near the ceiling. We use a fast, approximate logarithm first, reverting to a more accurate (and expensive) logarithm only if the first result is close to *K*_max_.

For fast logarithms, we use an approximation computed using the floating-point number’s integral representation. Specifically, we use a modified version of the algorithm of [18]. This can overestimate the result by a multiplicative factor of up to 1.42. By dividing the fast-logarithm result by this number, we can determine if the approximation is close enough to *K*_max_ to require a full logarithm computation. This affects the computation within RNG.nextExponentialSpacing(), called on lines 2 and 11, as well as the conditional checks on lines 4 and 12 of Algorithm 1, though we omitted these details from the algorithm listing.

### 2.8 Weighted SetSketch

While [8] describes applications only to unweighted sketching, we extended SetSketch to consider weightedness using the ProbMinHash [10] strategy, i.e. by multiplying the exponential draw’s rate parameter by the item’s weight, represented by argument *W* to RNG.nextExponentialSpacing() on lines 2 and 11 of Algorithm 1. When comparing sketches weighted in this way, the quantity being estimated is a version of the Jaccard coefficient called the “Probability Jaccard similarity” *J*_*P*_ :

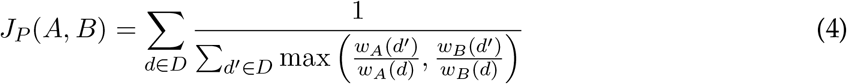

Where *D* is the item universe and *w*_*A*_ and *w*_*B*_ are weight functions for items in sets *A* and *B*.

A key question is how to obtain *w*_*A*_ and *w*_*B*_. When the input is sequencing data, the weight of an item (*k*-mer) should equal its relative frequency. By default, Dashing 2 will use a hash table to track the exact relative frequency for each item. But Dashing 2 also supports a faster and more memory-efficient method that estimates each item’s frequency using a feature hashing [19] approach, equivalent to a single-row Count-Min Sketch [20]. This mode is enabled with the `--countsketch-size` option. Non-sequencing datasets might also come with an inherent notion of “weight;” for instance, if input items represent genes and associated expression levels, these levels could be immediately used as weights, without the need for counting or for the feature-hashing data structure.

Optimizations for logarithms described above are also used for weighted sketching.

### 2.9 Locality-Sensitive Hashing (LSH) implementation

To scale all-pairs comparisons, we implemented a filtering approach based on locality-sensitive hashing (LSH). Given a minimum similarity threshold, the filter avoids computing many below-threshold pairwise comparisons. The LSH method works by grouping SetSketch registers into “super-registers.” For instance, the first four registers (*K*[0…3]) might constitute the first super-register, the next four (*K*[4…7]) the second super-register, etc. Associated with each super-register is a map from possible values to a list of all input datasets having that value in that super-register. In our example, the keys for the first super-register will consist of all combinations of the first four registers *K*[0…3] observed in an input dataset, and the values will be the associated lists of datasets. An LSH index might consist of several such tables, each with a distinct super-register group size. Algorithm S1 shows how the index is updated with one additional dataset.

Since registers are independent, a size-*P* super-register will match between datasets *A* and *B* with probability *J* (*A, B*)^*P*^, where *J* is the Jaccard coefficient. When performing a large-scale all-pairs comparison, Dashing 2 begins by computing LSH indexes for values of *P* in a user-configurable subset of the values {1, 2, 4, 6, 8, 10}. By default, Dashing 2 tries *P* ∈ {1, 2}, but it can be configured to try more values for *P* via the `--nlsh` option, corresponding to the *N* variable in Algorithms S1 and 2. For *P* ∈ {1, 2}, the super-registers are formed by partitioning registers into *m/P* non-overlapping groups. For larger values of *P*, we select a random set of *m*·8*/P* contiguous groups of registers. In this case, super-registers can overlap.

Algorithm S1 details how a single dataset is added to an LSH index. Algorithm 2 details how we query to find a list of candidate nearest neighbors for a dataset using the LSH index. In both cases, a pair of nested loops is used. The outer loop iterates over LSH tables from the most to least specific (largest to smallest *P*), while the inner loop iterates over super-registers. In the case of Algorithm S1, an iteration of the inner loop updates the LSH table with the *id* of the current dataset. In the case of Algorithm 2, an iteration of the inner loop contains a final loop that updates a running list of candidate datasets with all other datasets having the same value for the current super-register.

The LSH tables are used in two distinct modes of Dashing 2. The mode activated with (`--topk`) builds a *k*-nearest neighbor (KNN) graph from the input datasets and follows the logic of Algorithm 2. For a given pivot genome, we use the LSH tables to generate a list of [*O*_*s*_ × *k*] candidates for each input genome, where *O*_*s*_ *>* 1 is an over-sampling rate, set to 3 by default. We then estimate the Jaccard similarity between the pivot and each of the candidates in the order they were discovered, keeping only the *k* with the greatest Jaccard coefficients. While this can result in some misreported neighbors, e.g. because a near neighbor happened not to coincide with the pivot in any super-register, this possibility is reduced both by over-sampling and by the order in which we attempt the LSH tables, i.e. from most to least specific.

In another mode, Dashing 2 reports pairwise distances between all pairs of genomes having similarity above some threshold (`--similarity-threshold X`). In this mode, there is no additional limit on the number of “neighbors” that might be reported for a genome. While querying the index, we maintain a heap of all neighbors with similarity above a given threshold.

### 2.10 Exact similarity mode

The ability to compute exact Jaccard coefficients is useful for evaluating Dashing 2’s estimates. Dashing 2 therefore implements two modes for exact computation of Jaccard coefficients. One uses sorted *k*-mer hash sets (`--set`); the other uses *k*-mer count dictionaries (`--countdict`).

### 2.11 Sketching sequencing reads

Dashing 2 can also sketch inputs consisting of sequencing reads, e.g. in FASTQ format. This involves the extra challenge of handling sequencing errors, since *k*-mers containing errors can be far more numerous than correct *k*-mers and so can dominate and bias similarity estimates. For this reason, methods for sketching sequencing-read inputs attempt to filter out *k*-mers containing sequencing errors prior to computing cardinality or similarity. Dashing 2 adapts the approach of Mash [1] for eliminating k-mers below a specific count threshold. For instance, if the target threshold is set to 2, Dashing 2’s SetSketch implementations (both one-permutation and full) maintain a dictionary of items seen fewer than 2 times so far. Once an item’s count reaches 2, it is added to the final SetSketch structure.

Dashing 2 can also use a downsampling approach (`--downsample <fraction>`) to randomly keep a specified fraction of the input *k*-mers. The decision to keep or suppress a *k*-mer is made independently for each *k*-mer, rahter than for each *distinct k*-mer. In this way, frequent *k*-mers – i.e. occurring more than 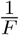 of the time – are unlikely to have all copies suppressed. The filter incurs little computational cost but greatly reduces the number of error *k*-mers ending up in the sketch. This is similar to the ideas used in sequencing error correction [21] to distinguish *k*-mers with or without sequencing errors.

Weighted sketching modes can be particularly appropriate for sequencing reads, since they have the effect of down-weighting error *k*-mers, which tend to occur infrequently compared to correct *k*-mers.

## 3 Results

We used Dashing 2 v2.1.11-10-g128c. All experiments were performed on a Lenovo ThinkSystem SR630 with 48 3.0GHz Xeon CPUs and 1.5 TB of memory.

We downloaded the Refseq database on Jun 30, 2022 [22]. Filtering to just complete genome sequences, we gathered 128,827 sequences, 729 from the “archea” category, 115,548 from “bacteria,” 338 from “fungi,” 2 from “human” (the GRCh38 and the CHM13 assemblies), 267 from “invertebrate,” 145 from “plant,” 88 from “protozoa,” 183 from “vertebrate mammalian,” 274 from “vertebrate other” and 11,253 from “viral.” The compressed FASTA files occupied 475 GB. Overall, genome lengths varied from 223 bp to over 34 billion bp, with mean and median lengths of 11.1 million and 4.22 million bp respectively.

Some experiments required high-fidelity sketches. We computed a 1-MB sketch for each using Dashing 2 with `sketch --binary-output -S 20 -k 31`. For all 128,827 complete genomes, this process took 50m:45s using GNU parallel [23] for multiprocess parallelism.

The following subsections use these assemblies as a starting point. In particular, Sections 3.1 and 3.1 & 3.2 use sets of 1,010 and 984 genome pairs selected to cover a range of similarities. Section 3.3 uses a subset of 50,000 assemblies to compare Dashing 2’s sketching and pairwise similarity speed to that of Dashing 1. The exact lists of accessions used in each experiment are provided in files referenced in the “Data and Software Availability” section.

### 3.1 Similarity estimation

We compiled a collection of pairs of assemblies covering a range of true Jaccard coefficients. If *A* and *B* are sets of canonicalized *k*-mers from two assemblies, the Jaccard coefficient *J* (*A, B*) = |*A* ∩ *B*|*/*|*A* ∪ *B*|. We first performed an all-pairs comparisons using the 128,827 high-fidelity sketches described above. We then partitioned the space of Jaccard estimates into 100 buckets of equal size. I.e. one bucket spanned Jaccard values in the range [0, 0.01), the next spanned values in (0.01, 0.02], etc. We added an additional bucket for pairs with *J* (*A, B*) = 1. For each bucket, we randomly selected 10 genome pairs having a Jaccard coefficient estimate within the bucket’s range. We limited our attention to Refseq assemblies from the “archea,” “bacteria” and “viral” groups. At the end of this process we had a collection of 1,010 genome pairs (10 for each of the 101 buckets) with Jaccard coefficients spread evenly across the range [0, 1].

To obtain a notion of “truth” to compare against, we used Dashing 2’s full-accuracy mode (which does not use sketching) to compute true Jaccard coefficients for all genome pairs for both *k*-mer lengths. We also computed Average Nucleotide Identities (ANIs) for all selected genome pairs using fastANI v1.33 [24]. fastANI computes an approximation, so we do not call these “true” ANIs. But these have the advantage of being calculated using a separate approach from the one used to compute the Jaccard coefficients; in particular, fastANI’s approach is not *k*-mer-based.

Using these genome pairs annotated with true Jaccard coefficients and fastANI-estimated ANIs, we compared the accuracy of Dashing 2’s estimates to those of Dashing 1 v1.0 [7] and Mash v2.3 [1]. We omitted BinDash initially as it would sometimes fail when comparing more distant genome pairs. We ran Dashing 2 in two configurations. **D2** used the “one-permutation” SetSketch, with each update affecting at most one register. **D2-full** used the full update rule. We did not run the **D2W** configuration of Dashing 2 here since the goal of these experiments is to assess how well these modes estimate the typical “flat” version of the Jaccard similarity, rather than the weighted version estimated by **D2W**.

Table 1 shows the sum of squared errors (SSE) between the tool-estimated Jaccard coefficient and the true Jaccard, totaled across all 1,010 genome pairs. We show these results for sketch sizes of 8 Kbits (8 × 1024 bits, equivalent to 1 Kbyte), 32 Kbits, and 128 Kbits. In all cases either D2 or D2-full achieved lowest SSE, with the other achieving second-lowest. Dashing 2 and Dashing 1 both achieved lower SSE than Mash.

**Table 1:** Sum of squared error between estimated and true Jaccard coefficients for several methods and for all genome pairs. Bright red indicates the lowest error in each row. Dark red indicates second-lowest error.

| k | kbits | Mash | Dashing1 | D2-full | D2 |
| --- | --- | --- | --- | --- | --- |
| 21 | 8 | 1.79 | 1.10 | 0.776 | 0.814 |
|  | 32 | 0.872 | 0.711 | 0.669 | 0.651 |
|  | 128 | 0.765 | 0.644 | 0.628 | 0.616 |
| 31 | 8 | 1.95 | 0.938 | 0.584 | 0.595 |
|  | 32 | 0.766 | 0.580 | 0.475 | 0.463 |
|  | 128 | 0.544 | 0.479 | 0.461 | 0.456 |

To additionally compare to BinDash, we filtered the 1,010 genome pairs down to the 984 pairs having fastANI-estimated ANI greater than or equal to 87%. For this subset, we were always able to run BinDash with no crashes. Table 2 shows these results for the same sketch sizes and values of *k* as Table 1. In all but one case, D2 achieved the lowest SSE; the exception was the case of *k* = 21 and 8 Kbit sketch, where BinDash achieved the lowest SSE. D2 or D2-full achieved the lowest SSEs in all other cases.

**Table 2:** Sum of squared error between estimated and true Jaccard coefficients for several methods, using only genome pairs with estimated true ANI greater than 89%. Above this threshold, BinDash ran with no estimation failures. Bright red indicates the lowest error and dark red the second-lowest error in each row.

| k | kbits | Mash | BinDash | Dashing1 | D2-full | D2 |
| --- | --- | --- | --- | --- | --- | --- |
| 21 | 8 | 1.78 | **0.750** | 1.08 | **0.773** | 0.812 |
|  | 32 | 0.870 | 0.676 | 0.705 | **0.668** | **0.651** |
|  | 128 | 0.764 | 0.648 | 0.642 | **0.628** | **0.615** |
| 31 | 8 | 1.94 | 0.600 | 0.912 | **0.583** | 0.594 |
|  | 32 | 0.764 | 0.480 | 0.573 | **0.475** | **0.463** |
|  | 128 | 0.544 | 0.473 | 0.477 | **0.461** | **0.456** |

### 3.2 ANI estimation

We further assessed the Average Nucleotide Identity (ANI) estimates obtained by using the Mash distance equation and re-scaling: *ANI*_est_ = 1 + 1*/k* · ln(2*J/*(1 + *J*)). Here *k* is the *k*-mer length and *J* is the estimated Jaccard coefficient. Negative *ANI*_est_ values were rounded up to 0. In this experiment, we additionally assessed the new multiplicity-aware (“weighted”) mode of Dashing 2, called **D2W**. The feature-hashing structure needed to obtain the weights for **D2W** mode was configured to consist of 5 million 64-bit counts.

In this case, the input *J* to the *ANI*_*est*_ equation was the probability Jaccard similarity (*J*_*P*_) described in section 2.8, rather than the typical “flat” Jaccard coefficient. This experiment allows us to assess whether *J* (as estimated by Dashing 1, D2 or D2-full) or *J*_*P*_ (as estimated by D2W) yields a better ANI estimate.

We assessed SSE between *ANI*_*est*_ and the fastANI-estimated ANIs for Mash, Dashing 1, D2 and D2W (Table 3). In all cases, the D2W approach achieved either the lowest or second-lowest SSE, with either D2 or Dashing 1 achieving the second-lowest SSE.

**Table 3:**
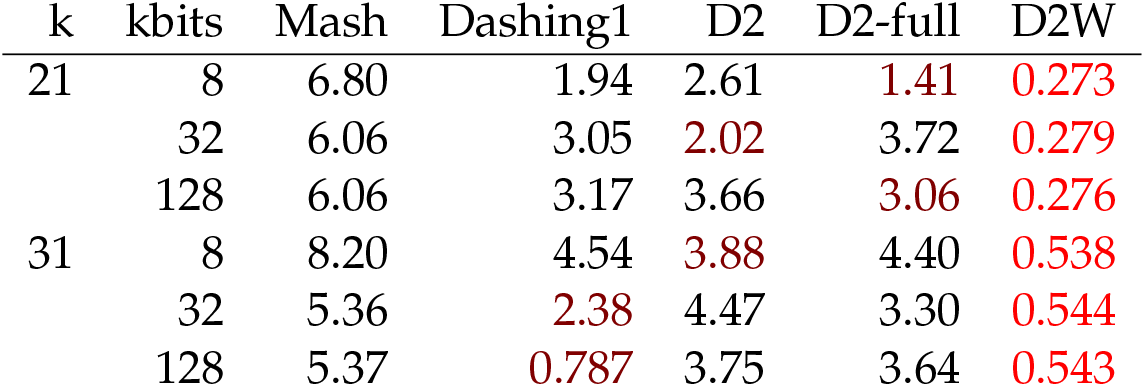
Sum of squared error between estimated Mash distance and the ANI as computed by fastANI. Bright red indicates the lowest error in each row. Dark red indicates second-lowest error. In all cases except D2W, Mash distance is computed as a function of the estimated “flat” Jaccard coefficient. In the case of D2W, Mash distance is a function of the Probability Jaccard Similarity from equation 4.

| k | kbits | Mash | Dashing1 | D2 | D2-full | D2W |
| --- | --- | --- | --- | --- | --- | --- |
| 21 | 8 | 6.80 | 1.94 | 2.61 | **1.41** | **0.273** |
|  | 32 | 6.06 | 3.05 | **2.02** | 3.72 | **0.279** |
|  | 128 | 6.06 | 3.17 | 3.66 | **3.06** | **0.276** |
| 31 | 8 | 8.20 | 4.54 | **3.88** | 4.40 | **0.538** |
|  | 32 | 5.36 | **2.38** | 4.47 | 3.30 | **0.544** |
|  | 128 | 5.37 | **0.787** | 3.75 | 3.64 | **0.543** |

### 3.3 Refseq sketching and pairwise comparisons

We used Dashing 1 and Dashing 2 to sketch a sample of 50,000 complete genome assemblies downloaded from Refseq. Both were run using GNU parallel, allowing up to 12 sketching processes to run at a time. Both tools were configured to produce a sketches 1 MB in size. Dashing 1 took 2h:00m:34s to construct all the sketches and Dashing 2 took 50m:46s.

We then used Dashing 1 and 2 to perform exhaustive all-pairwise Jaccard similarity comparisons across the 50,000 genomes (2 human assemblies and 49,998 bacterial assemblies), comparing a total of 1.25 billion pairs of 1MB sketches. Both tools used their default similarity estimation methods. In the case of Dashing 1’s, this was the MLE estimator of Ertl [13]. In the case of Dashing 2, this was the simple joint estimator described in section 2.2. Both Dashing 1 and Dashing were run with the `-p80 --presketched` options, enabling 80 simultaneous threads of execution and instructing both tools to use the already-computed sketches. Dashing 1 took 49h:41m to estimate all-pairwise similarities, whereas Dashing 2 took 5h:41m and was about 8.7 times faster.

### 3.4 All-pairs comparisons using LSH

We performed all-pairs comparisons for a large collection of proteins by combining SetSketch with locality-sensitive hashing to avoid comparisons unlikely to meet a minimum similarity threshold. We used the UniProtKB/Swiss-Prot collection of 565,254 protein sequences, v2021 03. For each protein, we used Dashing 2 to create a 10-mer sketch (`-k10`) of 256 registers (`-S256`). Proteins were translated to a 14-letter reduced alphabet (`--protein14`) to capture more distant homology [11]. We ran Dashing 2 in `sketch --topk` mode to perform all-pairs comparisons while avoiding pairings that fail to appear in the top 256 neighbors of a protein. We used the default of `--nLSH 2` to use two distinct sizes of super-register groupings, corresponding to the *N* parameter of Algorithm 2. The output was a K-Nearest-Neighbor (KNN) graph in tabular (TSV) format, associating each protein to the 256 others with greatest Jaccard coefficient.

Because of both estimation and LSH error, the graph may include false positives (reported neighbors that are not truly among the top 256) and/or false negatives (unreported neighbors that are truly in the top 256). To measure the error introduced by LSH, we compared the KNN graph to another KNN generated using exhaustive all-pairs comparisons. Importantly, the exhaustive KNN was also built using Jaccard coefficient estimates, since exact computation of the Jaccard coefficient is too computationally expensive. Thus, this experiment isolates error due to the LSH filter only, and does not assess error due to the Jaccard estimate.

Relative to the exact method, Dashing 2’s LSH method achieved 100% recall and precision, i.e. there were no false positives or false negatives among the neighbors found using LSH filtering. Further, using the LSH, Dashing 2 was able to generate the graph in 43 seconds, compared to 56.2 minutes for the exhaustive method, a 78-fold speedup. The exhaustive method ultimately performed 159,755,759,631 pairwise comparisons, compared to approximately 470M comparisons performed by the LSH-assisted method. The 470M comparisons performed by LSH is about 3.5 times greater than the minimum determined by the number of neighbors (256) times the number of proteins (565,254).

We also applied this approach to create a nearest-neighbor graph over the larger UniRef50 dataset, which is built by clustering UniRef90 seed sequences from the UniProt Knowledgebase v2021 03 having at least 50% sequence identity to and 80% overlap with the longest sequence in the cluster. The database contains 53,625,855 sequences totaling over 15 billion amino acids. After sketching, the KNN graph was generated in less than 10 minutes. By contrast, the exhaustive all-pairs comparisons approach required a much longer amount of time; we interrupted this computation after 2 days and extrapolated that the total time would have exceeded a year.

## 4 Discussion

Dashing 2 combines the SetSketch with locality-sensitive hashing to bring multiplicity-aware sketching and fast filtering to genomic sketching analysis. Its all-pairs comparison method scales effectively to millions of sequences. Dashing 2 can sketch FASTA and FASTQ inputs, as well as protein-sequence inputs using a variety of protein alphabet reductions. It can compare sequences based on Jaccard coefficient, Mash distance, or containment coefficient. It can also compare sequences based on a weighted version of the Jaccard coefficient that is aware of the multiplicities of the input items. Dashing 2 is free, open source software available at https://github.com/dnbaker/dashing2.

Dashing 2’s estimation error for the Jaccard coefficient and ANI estimates was lower than that of the previous version of Dashing, and substantially lower than that of Mash. In cases where BinDash can be run successfully, Dashing 2 had lower or comparable error to BinDash. Thus, it will be important to continue to study the SetSketch as a highly accurate alternative not only to the HyperLogLog sketch, but also the MinHash and b-bit MinHash methods.

It is remarkable that Dashing 2’s ANI estimation error was lowest when taking multiplicities of the input items into account, i.e. when using a weighted Jaccard coefficient as input to the Mash distance equation. This demonstrates that multiplicities are helpful not only for quantitative applications, but also in typical “flat” sketching scenarios. This motivates future study of sketch data structures that account for multiplicities, as well as methods – like Dashing 2’s feature hashing method – for efficiently compiling multiplicity information prior to sketching.

While the term “sketch” is used for various data structures in Bioinformatics, Dashing 2 is designed for the scenario where many sequencing datasets must be compared to each other in an efficient, scalable way. In these scenarios, we use the expedient of modeling the data as a set (or multiset) of *k*-mers. Other studies describe sketching approaches that keep more detail about the sequence content of each dataset, e.g. by keeping a ordered sequence of representative sub-sequences from the longer sequence [25, 26]. The OrderMinHash study further points out that a sketch for edit distance can be improved by taking *k*-mer multiplicity into account, a similar observation to the one we make in Table 3. But the advantages of such sketches come at the expense of slower algorithms and larger memory footprint; in particular, these sketches may grow linearly with the length of the sequence being sketched. That said, applications requiring more detailed knowledge of the sequences, e.g. when distinguishing between similar strains or identifying particular alleles of interest, can benefit from these more detailed sketches.

## 5 Acknowledgements

We thank Otmar Ertl and Martin Steinneger for helpful conversations. This work used the Extreme Science and Engineering Discovery Environment (XSEDE), which is supported by National Science Foundation grant number ACI-1548562. In particular, we acknowledge Texas Advanced Computing Center (TACC) at The University of Texas at Austin for providing HPC resources that have contributed to the research results reported within this paper. URL: http://www.tacc.utexas.edu.

## 7 Author contributions

DNB and BL conceived the study, designed the experiments, wrote the experimental scripts, ran the experiments and wrote the manuscript. DNB wrote the Dashing 2 software.

## 8 Data and Software Availability

Open source source code for the Dashing 2 software is at https://github.com/dnbaker/dashing2

Scripts for performing the experiments described in this manuscript are at https://github.com/dnbaker/dashing2-experiments

A list of the accessions for the 1,010 Refseq genome pairs used in Results 3.1 is at https://www.cs.jhu.edu/~langmea/resources/d2/pairs1010.csv

Accessions for the 984 Refseq genome pairs used in Results 3.2 are at https://www.cs.jhu.edu/~langmea/resources/d2/pairs984.csv

Accessions for the 50,000 Refseq genomes used in the experiments in Results 3.3 are at https://www.cs.jhu.edu/~langmea/resources/d2/refseq50k.txt.

## Footnotes

1 This estimator is described only in the “v1” version of the paper pre-print cited. Later versions of the paper describe Brent’s root-finding algorithm instead.

