## Supplementary Information for "Dashing 2: genomic sketching with multiplicities and locality-sensitive hashing"

### Supplementary Material

See following pages.

**Input:** $K[0..m-1]$: SetSketch registers for this dataset
**Input:** $id$: Identifier for this dataset
**Input:** $RNG$: Pseudo-random number generator
**Input:** $seed$: Pseudo-random seed
**Input:** $N$: Number of super-register sizes in index
**Input:** $LSH$: map from ⟨ table id, super-register id, super-register value ⟩ triples to a corresponding list of datasets
**Result:** $LSH$ updated to include dataset $\langle K, id \rangle$

i ← $N - 1$
**if** $i > 2$ **then**
    $RNG$.initialize($seed$)
**end**
// Loop over tables 0 .. $N$-1, from largest super-register size (most specific)
// to smallest (least specific)
**while** $i \geq 0$ **do**
    $P \leftarrow \min(2^i, 2i)$
    **if** $i \leq 2$ **then**
        S ← $m/N$
    **else**
        S ← $m \cdot 8/N$
    **end**
    $j \leftarrow 0$ // Loop over super-registers
    **while** $j < S$ **do**
        **if** $i \leq 2$ **then**
            // Next non-overlapping super-register
            SuperReg ← $K[P \cdot j \; .. \; P \cdot j + P - 1]$
        **else**
            // Get uniform random integer in [0, $m - P$]
            $ri$ ← RNG.randomInt(0, $m - P$)
            SuperReg ← $K[ri \; .. \; ri + P - 1]$
        **end**
        // Append this dataset to the list for this table, super-register
        // super-register value combination
        $LSH[\langle i, j,$ SuperReg $\rangle]$.append($id$)
        $j \leftarrow j + 1$
    **end**
    $i \leftarrow i - 1$
**end**

**Algorithm S1:** Add dataset $\langle K, id \rangle$ to LSH index

**Input:** SetSketch $K[0..m-1]$, item $X$
**Result:** $K$ updated according to $X$

$\text{RI} \leftarrow \text{hash}(X)$
$q \leftarrow \text{RI} \mod m$
$p \leftarrow \lfloor \text{RI}/m \rfloor$
$K[i] \leftarrow \min(K[i], p)$

**Algorithm S2:** Update one-permutation Dashing 2 SetSketch. Each $K_i$ gets the minimal 64-bit random draw among draws that map there.

**Input** : SetSketch $K[0..m-1]$, item $X$
**Output:** $K'$ set to a truncated form of $K$

**for** $i \leftarrow 0, m-1$ **do**
  $K'[i] \leftarrow \lfloor 1 - \log_b(K[i]) \rfloor$
**end**

**Algorithm S3:** Finalize Full Dashing 2 SetSketch

**Input** : Temporary $K[0..m-1]$ from Algorithm S2
**Output:** Finalized $K'$

**for** $i \leftarrow 0, m-1$ **do**
  $RV \leftarrow -\ln(K[i])$
  $K'[i] \leftarrow \lfloor 1 - \log_b(RV) \rfloor$
**end**

**Algorithm S4:** Finalize one-permutation Dashing 2 SetSketch
